# Plant-parasitic nematodes produce functional mimics of plant PSK peptides to facilitate parasitism

**DOI:** 10.64898/2026.04.04.713059

**Authors:** Yali Zhang, Dadong Dai, Vera Putker, Lena M. Müller, Sagar Bashyal, Shahid Siddique

## Abstract

Root-knot nematodes are obligate plant parasites that cause substantial agricultural losses worldwide. They induce highly specialized, metabolically hyperactive feeding sites within host roots, which serve as their sole source of nutrients throughout their life cycle. The formation and maintenance of these feeding sites depend on the manipulation of host developmental pathways by nematode-derived secretions. Phytosulfokines (PSKs) are small plant peptide hormones that regulate cell division, tissue expansion, and growth responses, processes essential for feeding site development. Here, we identify root-knot nematode genes predicted to encode peptides with a conserved PSK functional motif. These genes are predominantly expressed during the early stages of infection and localize to secretory glands, suggesting a role in early parasitism. Moreover, silencing PSK-like gene expression reduces root gall formation and nematode reproduction. Together, these findings reveal that root-knot nematodes deploy PSK-like peptides as virulence factors to promote successful parasitism, providing the first report of PSK peptide mimicry in any plant pathogen.

## Introduction

Plant-parasitic nematodes (PPNs) are a highly diverse group of obligate parasites with exceptionally broad host ranges. To date, more than 4,100 PPN species have been described, collectively infecting nearly all cultivated plants and causing annual global agricultural losses estimated at $157-173 billion (Zhu et al., 2022; Singh et al., 2015; Desmedt et al., 2020). Among them, root-knot nematodes (RKNs; *Meloidogyne* spp.) represent the most polyphagous and destructive genus worldwide (Jones et al., 2013). RKNs infect more than 2,000 plant species, including both monocotyledonous and dicotyledonous hosts. Although over 90 RKN species have been described, *M. incognita*, *M. arenaria*, *M. javanica*, and *M. hapla* are the most economically important (Rutter et al., 2022; Blundell et al., 2026).

Following root invasion, second-stage juveniles of RKNs (J2s) migrate to the vascular cylinder, where they induce the formation of five to seven highly specialized feeding cells known as giant cells (GCs). These giant cells serve as the nematode’s sole source of nutrients throughout its approximately one-month-long life cycle (Bartlem et al., 2013; Lin and Siddique, 2024). The establishment and long-term maintenance of giant cells are accompanied by extensive developmental reprogramming of surrounding tissues, resulting in localized cell proliferation and tissue expansion that ultimately give rise to characteristic root galls (Siddique and Grundler, 2018).

To establish and maintain these specialized feeding sites, RKNs secrete a wide range of molecules into host tissues, including proteins, hormones, and small peptides that manipulate host cellular processes (Bali and Gleason, 2024). Manipulation of host development through molecular mimicry is a common strategy among plant-associated pathogens. In PPNs, this strategy is most clearly exemplified by the secretion of small peptides that resemble endogenous plant signaling ligands, termed peptide mimics (Gheysen and Mitchum, 2019; Mitchum and Liu, 2022). The best-characterized peptide mimics are CLE (CLAVATA3/Embryo Surrounding Region-related) peptides. Following delivery into the host cytoplasm, CLE peptides are processed by host post-translational machinery into the mature, bioactive form before being released into the apoplast (Gheysen and Mitchum, 2019; Guo et al., 2011; Wang et al., 2021; Mitchum and Liu, 2022). Once in the apoplast, CLE mimics are perceived by membrane-localized CLAVATA receptor complexes, leading to host cell identity reprogramming and promotion of feeding site formation (Gheysen and Mitchum, 2019).

Beyond CLEs, nematodes also encode mimics of several other plant peptide families, including CEP (C-terminally encoded peptides), IDA/IDL (Inflorescence Deficient in Abscission peptides), and RALF (Rapid Alkalization Factor). These mimics manipulate the host nitrogen signaling, cell separation process, and immune responses, respectively (Zhang et al., 2020; Bobay et al., 2013; Kim et al., 2018). More recently, RKNs were shown to encode sulfated peptides with sequence and functional similarity to plant PSY peptides (Plant peptide containing sulfated tyrosine), whose biological activity depends on tyrosine sulfation (Yimer et al., 2023). These nematode-derived PSY-like peptides promote root growth and are required for successful parasitism, suggesting that nematodes can exploit host PSY signaling pathways during infection.

Phytosulfokines (PSKs) are highly conserved, tyrosine-sulfated pentapeptides that regulate cell division and expansion, meristem maintenance, immune signaling, and responses to abiotic stress (Li et al., 2024). PSKs are synthesized as precursor proteins of approximately 80–120 amino acids that contain an N-terminal signal peptide for entry into the secretory pathway, a variable region, and a conserved PSK functional motif near the C-terminus (Sauter, 2015). These precursors undergo post-translational processing, including proteolytic cleavage and tyrosine sulfation, to generate mature, biologically active pentapeptides (Stuhrwohldt et al., 2021; Reichardt et al., 2020; Sauter, 2015). Five distinct types of functional PSK peptides have been described so far: PSK-alpha (PSK-α), PSK-beta (PSK-β), PSK-gamma (PSK-γ), PSK-delta (PSK-δ), and PSK-epsilon (PSK-ε), which all share a conserved sulfation-dependent core motif. The canonical form, PSK-α (Y_SO3_IY_SO3_TQ), exhibits the highest biological activity (Matsubayashi et al., 1996). PSK-β (Y_SO3_IY_SO3_T) lacks the C-terminal glutamine, shows greatly reduced activity, and is therefore considered a degradation product of PSK-α (Yang et al., 1999). Other PSK variants differ mainly at the second and fifth amino acid positions: PSK-γ has an isoleucine-to-valine substitution at position 2 (Y_SO3_VY_SO3_TQ), PSK-δ has a glutamine-to-asparagine substitution at position 5 (Y_SO3_IY_SO3_TN), and PSK-ε contains both substitutions (Y_SO3_VY_SO3_TN)(Yu et al., 2019; Yu et al., 2022; Di et al., 2022). Tyrosine sulfation is catalyzed by tyrosylprotein sulfotransferases (TPSTs) and is essential for full PSK activity, as non-sulfated PSK exhibits strongly reduced biological function (Kutschmar et al., 2009). Consistent with this requirement, loss of TPST function in Arabidopsis results in dwarfism and root growth defects, underscoring the importance of PSK and related sulfated peptides in plant development (Komori et al., 2009). Active PSKs are perceived by membrane-localized PSK receptors (PSKRs), members of the leucine-rich receptor-like kinase family (Sauter, 2015). In both Arabidopsis and tomato, two PSKRs have been identified and functionally characterized to regulate growth and developmental responses (Matsubayashi et al., 2006; Stuhrwohldt et al., 2015; Amano et al., 2007; Zhang et al., 2018).

Despite the extensive knowledge of PSK biosynthesis, perception, and downstream signaling in plant developmental systems, no pathogen-derived PSK mimics have been reported to date. This absence is notable given that PSK signaling regulates cell division and tissue expansion, two mechanisms exploited by root-knot nematodes to facilitate feeding site formation. Additionally, loss-of-function mutations in PSK receptors in Arabidopsis have been shown to impair root-knot nematode infection, resulting in reduced host susceptibility and compromised nematode development (Rodiuc et al., 2016). Here, we identify root-knot nematode genes encoding PSK-like peptides with a conserved functional motif. We further characterize their expression and demonstrate their contribution to RKN parasitism.

## Results

### Plant-parasitic nematode genes encode PSK-like precursor proteins

To identify PSK candidates within PPN genomes, we implemented a comprehensive bioinformatics search integrating Hidden Markov Models (HMMER) and BLAST-based searches. We initially created a search profile using full-length PSK protein sequences from *Arabidopsis thaliana* and *Glycine max* (**Supplementary Table 1**). The inclusion of *G. max* ensured the representation of various PSK classes within the profile species (Li et al., 2024). This profile was then used to perform an initial HMMER search against the *Meloidogyne enterolobii* proteome available via the UniProt and WormBase ParaSite databases. After the initial identification of PSK sequences in PPNs, we used PSK sequences from RKNs as queries to perform BLAST searches against all available PPN protein databases (Blanc-Mathieu et al., 2017; Howe et al., 2017; Dai et al., 2023; Dai et al., 2026b). In parallel, we conducted an expression search to identify all protein sequences containing the DY*YT* motif. To ensure the selection of high-confidence candidates, sequences were required to meet four criteria: (i) presence of the DY*YT* motif; (ii) protein length <120 amino acids; (iii) presence of a signal peptide; and (iv) absence of transmembrane domains. By cross-validating the results from both approaches, we identified a total of 33 candidate protein sequences harboring the PSK motif across closely related root-knot nematode species belonging to *Meloidogyne* clade I, including *M. incognita*, *M. javanica*, *M. arenaria*, *M. floridensis*, and *M. enterolobii*. Beyond the root-knot nematodes, PSK-like sequences were also identified in soybean cyst nematode *Heterodera glycines* and the lesion nematode *Pratylenchus vulnus* (**Supplementary Table 1**). All identified nematode PSK-like sequences were encoded within short proteins with a predicted N-terminal secretion signal, followed by a propeptide region and a C-terminal PSK-like motif (**Supplementary Table 1**). Given the notably higher prevalence of PSK-like candidates in root-knot nematodes, we focused subsequent structural and evolutionary analyses on the RKNs.

Full-length sequence analysis of RKNs PSK-like peptides revealed strong conservation of hydrophobic residues in N-terminal signal peptide, substantial conservation within the propeptide region, and a highly conserved PSK-like motif at the C terminus (**Fig. 1A**).

**Figure 1.**
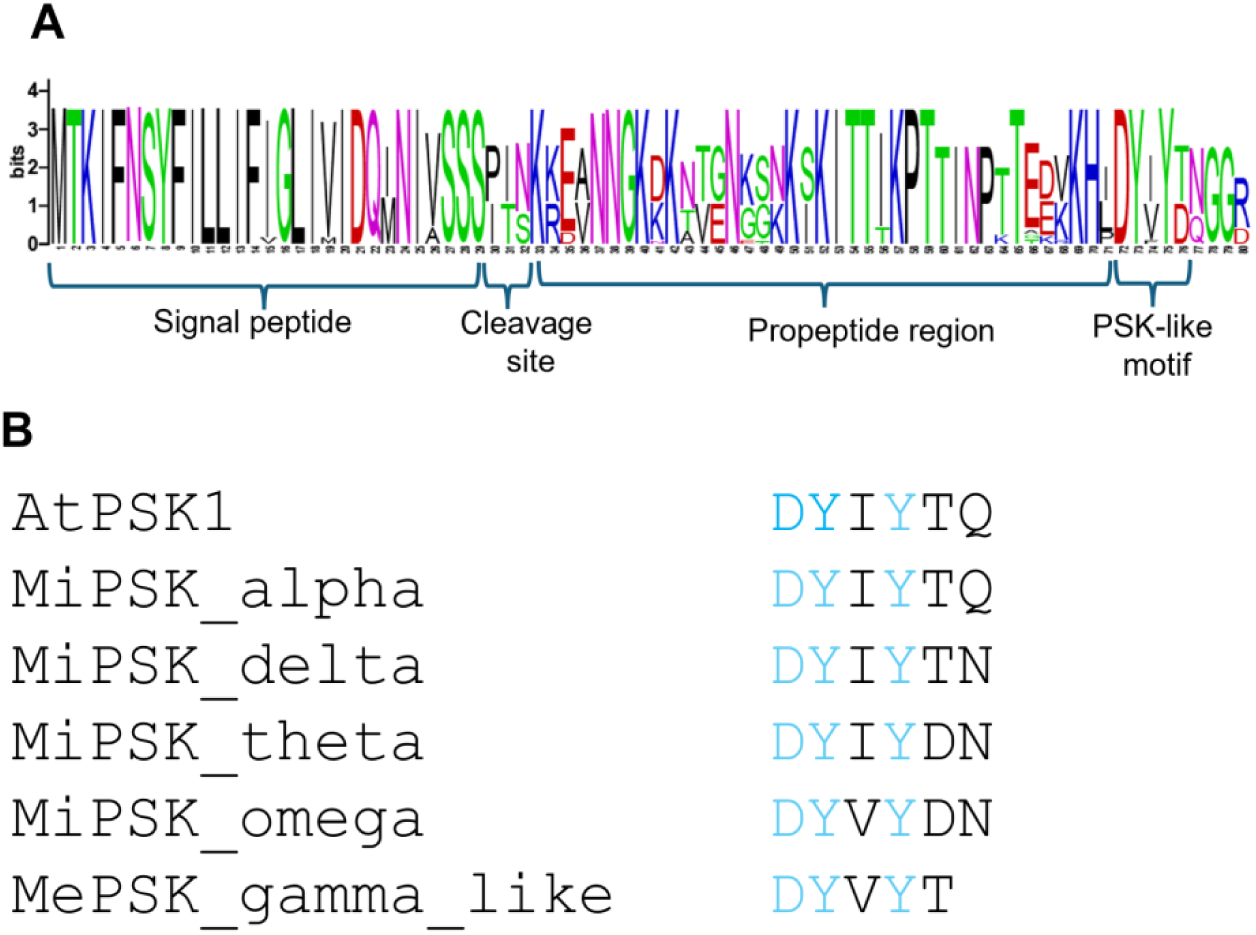
Root-knot nematodes encode peptides with a conserved C-terminal motif. (A) Protein sequence logo alignment of predicted PSK-like precursor proteins from root-knot nematodes. The alignment reveals a conserved N-terminal signal peptide, followed by a signal peptide cleavage site, a variable propeptide region, and a conserved C-terminal PSK-like motif. (B) Amino acid sequences of the conserved PSK motif from *Arabidopsis thaliana* AtPSK1 (DYIYTQ) compared with predicted PSK-like motifs identified in root-knot nematodes. Representative motifs from the five PSK-like classes are shown: MiPSK-alpha (DYIYTQ), MiPSK-delta (DYIYTN), MiPSK-theta (DYIYDN), MiPSK-omega (DYVYDN), and MePSK-gamma-like (DYVYT). The conserved DY sulfation site and the second tyrosine residue are indicated in blue. Variations at positions three, five, and six distinguish the different PSK-like classes.

Based on conserved amino acid substitutions within the C-terminal PSK motif, nematode PSK-like peptides were grouped into five classes: alpha (α), delta (δ), theta (θ), omega (ω), and gamma-like (γ-like) (**Table 1**; **Supplementary Table 1**). All classes share the invariant DY sulfation consensus required for TPST-mediated tyrosine sulfation and differ primarily at positions 3, 5, and 6 of the core motif. The α class (DYIYTQ) is identical to the canonical plant PSK-α motif and the most widely distributed, present across all examined *Meloidogyne* clade I species and *Pratylenchus vulnus*. The δ class (DYIYTN) differs from α at position 6, where glutamine is replaced by asparagine (Q→N), analogous to the plant PSK-δ isoform (Yu et al., 2022). The θ class (DYIYDN) carries an additional threonine-to-aspartate substitution at position 5 (T→D) not found in any plant PSK, making it a nematode-specific variant. The ω class (DYVYDN) has valine at position 3 (I→V) and terminates in a conserved GGR tripeptide, a propeptide amidation signal characteristic of nematode neuropeptide precursors (Li, 2018; Rosoff et al., 1993). The γ-like class (DYVYT) is a five-residue motif resembling plant PSK-γ but lacking the sixth terminal residue. Beyond these five classes, two divergent subclasses were identified in subsets of species: an ω-like variant (EVFYDN) in *M. javanica* and *M. arenaria*, and α-like sequences in *Heterodera glycines* (DYIYTE), which differ from the canonical α motif only by a glutamate at the terminal position (Q→E). Notably, in RKNs, multiple PSK-encoding sequences were clustered on the same chromosome **(Supplementary Fig. 1)**. This clustering may facilitate coordinated transcription or ensure robust production of PSK-like peptides during parasitic stages, potentially enhancing their effectiveness for host manipulations.

**Table 1:** Classification and sequence features of nematode PSK-like peptide classes. PSK-like peptides identified in plant-parasitic nematodes were grouped into five distinct classes based on conserved amino acid substitutions within the PSK core motif. The canonical PSK motif (DYIYTQ) defines the α class, whereas the remaining classes display substitutions at specific positions within the motif and/or variation in the terminal residue. Mi, *Meloidogyne incognita*; Mf, *M. floridensis*; Me, *M. enterolobii*; Ma, *M. arenaria*; Mj, *M. javanica*; Pv, *Pratylenchus vulnus*; Hg, *Heterodera glycines*.

| PSK-like class | Core motif | Closest plant class | Nematode Species |
| --- | --- | --- | --- |
| Alpha (α) | DYIYTQ | PSK-α (Li et al., 2024) | Ma, Me, Mf, Mi, Mj, Pv, Hg |
| Gamma-like (γ-like) | DYVYT | PSK-γ (Li et al., 2024) | Ma, Me, Mj |
| Delta (δ) | DYIYTN | PSK-δ (Yu et al., 2022) | Ma, Me, Mi, Mj, |
| Theta (θ) | DYIYDN | None (nematode-specific) | Ma, Me, Mi, |
| Omega (ω) | DYVYDN | None (nematode-specific ) | Ma, Me, Mf, Mi, Mj |

### Phylogenetic analysis reveals lineage-specific expansion of nematode PSK-like peptides

To investigate evolutionary origins of nematode PSK-like proteins, we performed a phylogenetic analysis using nematode full length PSK-like protein sequences alongside representative plant PSK proteins from Arabidopsis, tomato, and soybean. Our results reveal a striking divergence between the nematode and plant PSKs, with PSK-like proteins from the root-knot nematodes forming a highly supported, monophyletic clade. This cohesive clustering suggests that nematode PSK-like genes may have undergone a lineage-specific expansion, potentially to facilitate specialized host parasite signaling. Interestingly, *H. glycines* formed a distinct clade as compared to the other RKN PSK-like sequences (Fig. 2). The PSK-like sequence from *P. vulnus* occupied a position basal to the root-knot nematode cluster, consistent with its phylogenetic placement relative to *Meloidogyne* species (Dai et al., 2026b). Taken together, these relationships suggest that while PSK-like peptides represent a conserved feature of plant parasitic nematodes, they have evolved into a diversified gene family from their host counterparts.

**Figure 2.**
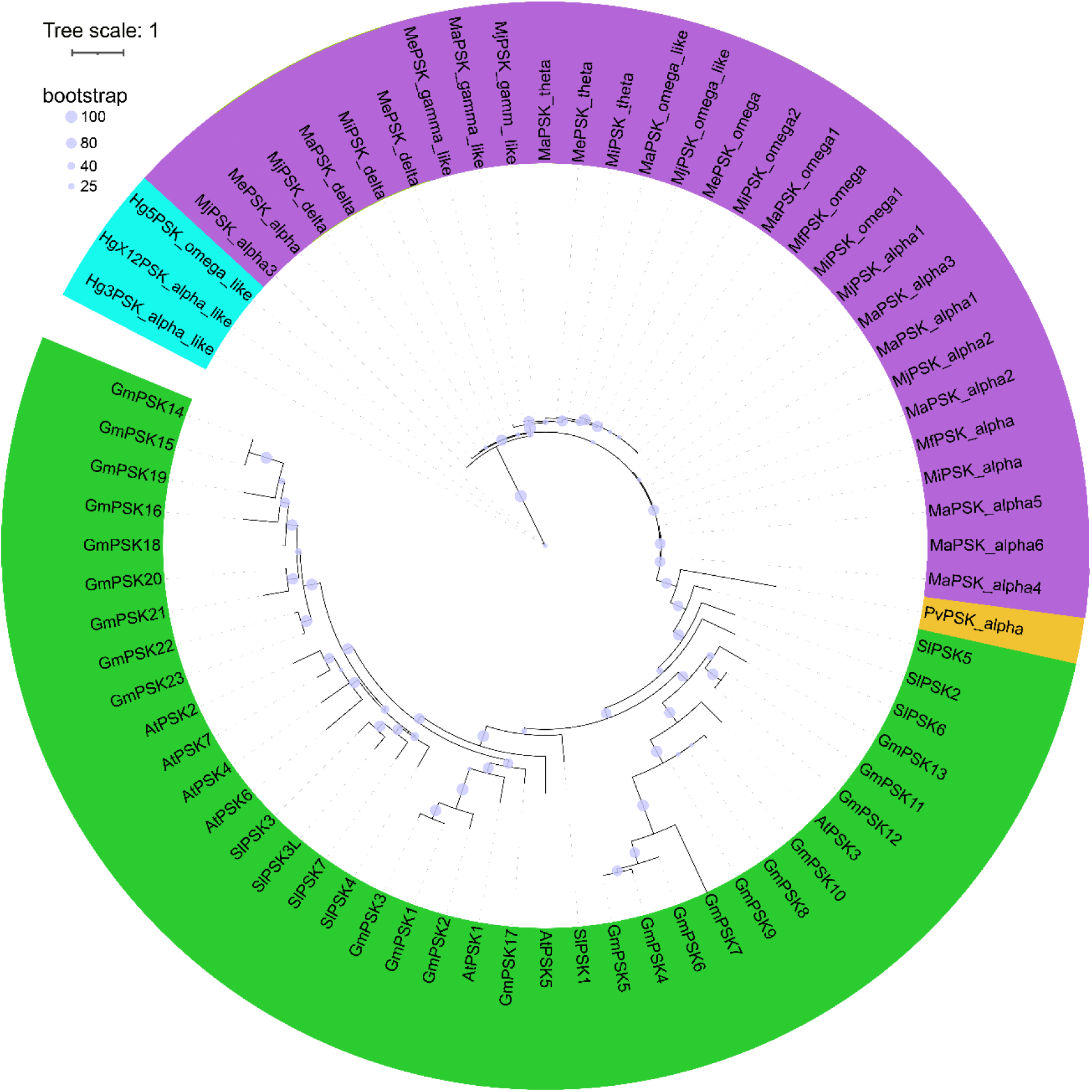
Phylogenetic analysis of plant and nematode PSK and PSK-like proteins. Maximum-likelihood phylogenetic tree constructed using predicted PSK-like proteins full-length sequence from plant-parasitic nematodes and representative plant species, including *Arabidopsis thaliana* (At), tomato (*Solanum lycopersicum*; Sl), and soybean (*Glycine max*; Gm). Plant PSKs are indicated in blue. Root-knot nematode (*Meloidogyne* spp.) PSK-like proteins are shown in green, the soybean cyst nematode (*Heterodera glycines*) sequences are highlighted in teal, and the lesion nematode (*Pratylenchus vulnus*) sequence is indicated in yellow. Bootstrap support values are represented by circles at the nodes, with size corresponding to support (25–100). The scale bar represents 1 substitution per site.

### *MiPSK* transcripts are expressed in the esophageal glands of nematodes

To assess the potential secretory nature of MiPSK peptides, we performed *in situ* hybridization to localize MiPSK transcripts in root-attracted J2s of *M. incognita*. Hybridization with a pooled mixture of DIG-labeled antisense probes derived from three *MiPSK* genes representing the α, θ, and ω classes revealed specific signals localized to the subventral esophageal glands (SvG; **Fig. 3A**), the principal secretory glands active during the infective J2 stage and the primary source of effector proteins delivered into host tissue via the stylet. An additional hybridization signal was detected in the posterior region of the nematode body (**Fig. 3C**). The identity of this structure remains uncertain — it may correspond to the phasmids, paired chemosensory organs in the tail region that open to the exterior and have been shown to detect chemical cues in the environment (Hilliard et al., 2002), or alternatively to the rectal glands, which have been less extensively characterized in the context of parasitism. Distinguishing between these possibilities will require gene-specific probes for each MiPSK class individually, as the pooled probe approach used here does not resolve which transcripts contribute to the posterior signal or whether this localization pattern is shared across all classes or restricted to a subset. No signal was observed with DIG-labeled sense probes (To assess the potential secretory nature of MiPSK peptides, we performed *in situ* hybridization to localize *MiPSK* transcripts in root-attracted J2s of *M. incognita*. Hybridization with a pooled mixture of DIG-labeled antisense probes derived from three *MiPSK* genes representing the α, θ, and ω classes revealed specific signals localized to the subventral esophageal glands (SvG; **Fig. 3A**), the principal secretory glands active during the infective J2 stage and the primary source of effector proteins delivered into host tissue via the stylet. An additional hybridization signal was detected in the posterior region of the nematode body (**Fig. 3C**). The identity of this structure remains uncertain, it may correspond to the phasmids, paired chemosensory organs in the tail region that open to the exterior and have been shown to detect chemical cues in the environment (Hilliard et al., 2002), or alternatively to the rectal glands, which have been less extensively characterized in the context of parasitism. Distinguishing between these possibilities will require gene-specific probes for each *MiPSK* class individually, as the pooled probe approach used here does not resolve which transcripts contribute to the posterior signal or whether this localization pattern is shared across all classes or restricted to a subset. No signal was observed with DIG-labeled sense probes (**Fig. 3B, D**). Taken together, these findings suggest that *MiPSK* peptides are produced in esophageal glands and possibly additional secretory tissues and are likely delivered into host cells during the early stages of infection.

**Figure 3.**
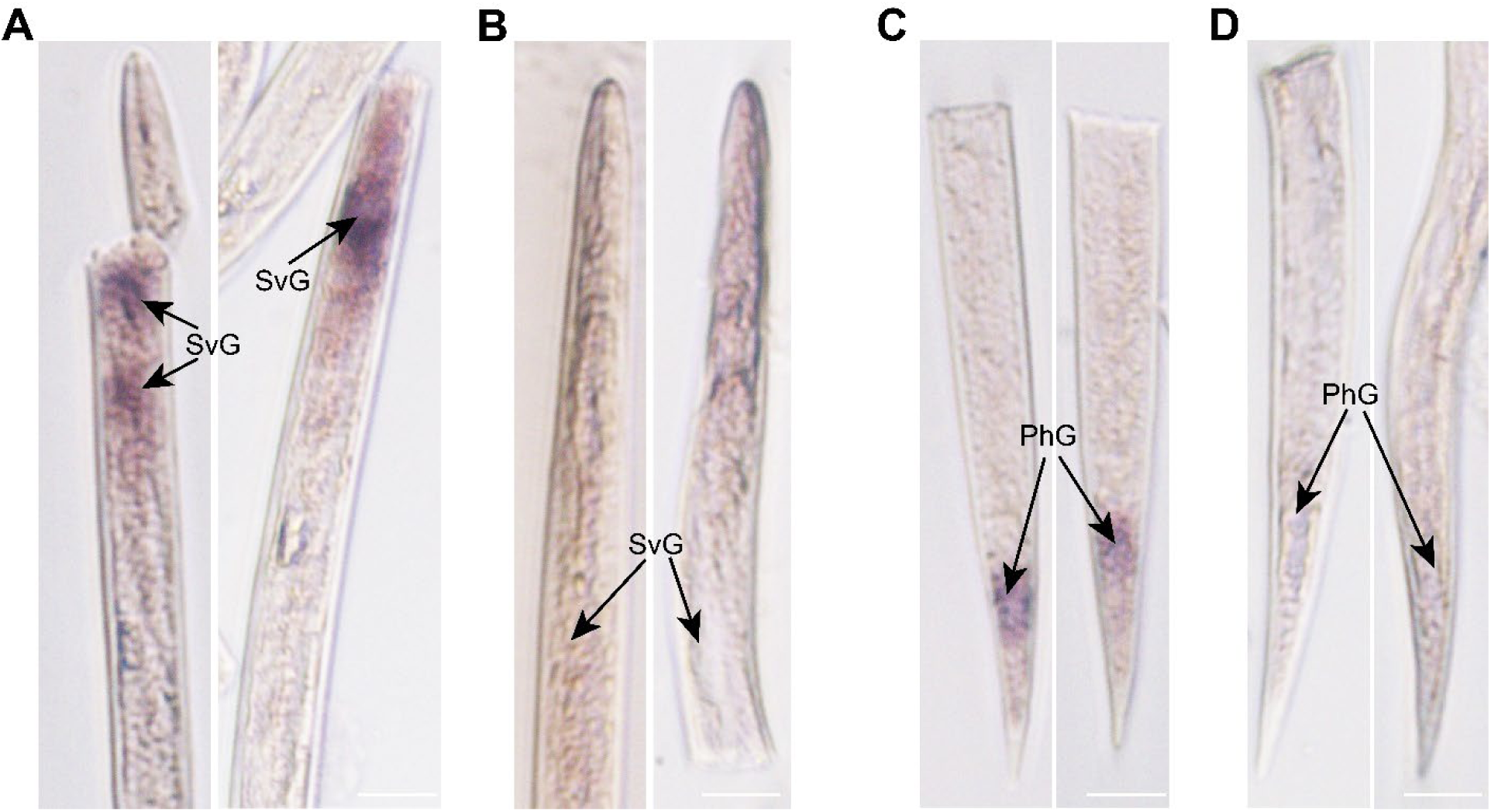
In situ hybridization of *MiPSK* transcripts in *M. incognita*. (A, C) Detection of MiPSK transcripts in root-attracted J2s using a pooled mixture of antisense DIG-labeled probes derived from *MiPSK-α*, *MiPSK-θ*, and *MiPSK-ω*. Hybridization signals were detected in the subventral esophageal glands (SvG; A) and in the posterior region of the nematode body (C), the identity of which — phasmid or rectal gland — remains to be determined. (B, D) Sense probes used as negative controls showed no detectable signal. DIG, digoxigenin; SvG, subventral esophageal glands. The experiment was performed three independent times with similar results. Representative images are shown. Scale bars = 20 μm.

### *MiPSK* gene expression is induced during early stages of nematode infection

To investigate the temporal regulation of *MiPSK* genes (*MiPSK-α*, *MiPSK-δ*, *MiPSK-θ*, and *MiPSK-*ω) during host recognition and parasitism, we performed quantitative real-time PCR (qRT-PCR) across different developmental stages. These included eggs, pre-parasitic J2s (pre-J2s), root-attracted J2s, sedentary parasitic stages (par-J3/J4s), and adult females. Root segments containing galls were collected from tomato plants inoculated with *M. incognita* to obtain eggs, par-J3/J4s, and adult females. Pre-J2s were obtained by hatching eggs isolated from infected tomato roots, whereas root-attracted J2s were collected after exposing tomato roots to *in vitro*–hatched J2s for 6 hours. Total RNA was extracted from nematodes at each developmental stage, and cDNA was synthesized and used as a template for qRT-PCR with gene-specific primers (supplemental table 2).

All four *MiPSK* genes exhibited strong expression in root-attracted J2s compared with other developmental stages (**Fig. 4**). In addition, *MiPSK-θ* and *MiPSK-*ω transcripts were significantly upregulated in pre-parasitic juveniles. In contrast, the expression levels of all *MiPSK* genes declined sharply during the sedentary parasitic stages and were barely detectable in adult females. This expression pattern closely resembles that reported for PSY peptide mimics and supports a role for PSK-like peptides during early parasitism, particularly in the initiation of feeding site (giant cell) formation (Yimer et al., 2023).

**Figure 4.**
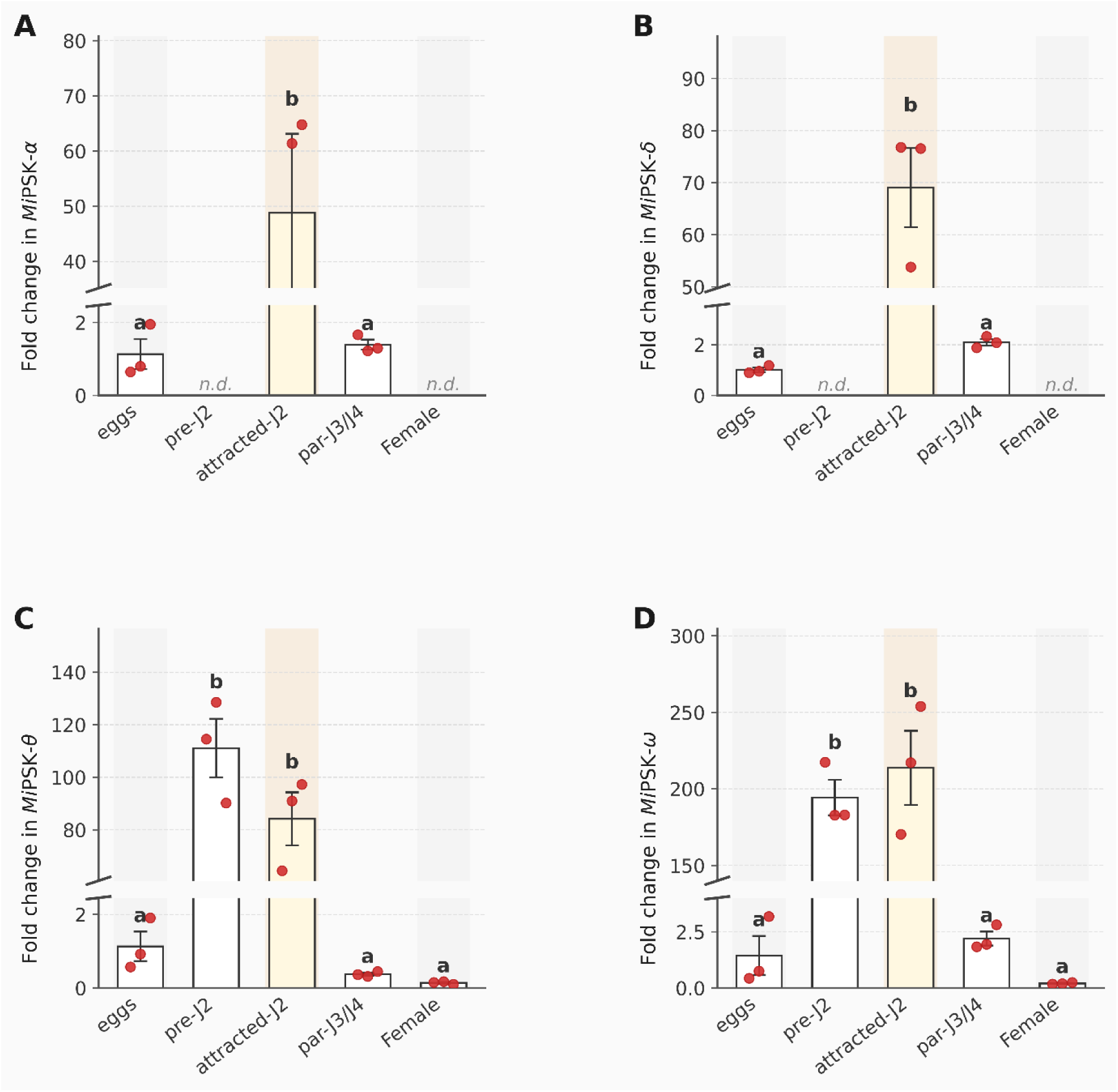
MiPSK genes are induced during the early stages of nematode infection. Relative expression levels of MiPSK-α (A), MiPSK-δ (B), MiPSK-θ (C), and MiPSK-ω (D) across five developmental stages of *Meloidogyne incognita*: eggs, pre-parasitic second-stage juveniles (pre-J2), root-attracted J2s (attracted-J2), sedentary parasitic juveniles (par-J3/J4), and adult females. Transcript levels were quantified by quantitative real-time PCR (qRT-PCR) and normalized to the housekeeping gene 18S rRNA using the 2^−ΔΔCt method (Livak and Schmittgen, 2001), with the egg stage set as the calibrator (fold change = 1). Expression in pre-J2 and adult female stages could not be reliably quantified for MiPSK-α and MiPSK-δ (n.d., not detected). Bars represent the mean ± SEM of three technical replicates. Red circles indicate individual data points. Statistical significance was assessed by one-way ANOVA followed by Dunnett’s multiple comparison test, with the egg stage as the reference group (set at 1). Different letters above bars indicate statistically significant differences (P ≤ 0.05). Data shown are from one representative biological replicate; a second independent experiment yielded consistent results. A third independent biological replicate is currently in progress and will be included in the final peer-reviewed manuscript.

### Silencing of *MiPSKs* compromises gall formation and parasitic success

To investigate if MiPSKs contribute to host infection, pre-parasitic J2s of *M. incognita* were soaked in specific double-stranded RNA (dsRNA) targeting *MiPSK-θ, MiPSK-ω* and *MiPSK-α*. Due to the close sequence similarity between MiPSK-α and MiPSK-δ, a single dsRNA construct was designed targeting MiPSK-α, which is expected to also silence MiPSK-δ through cross-reactivity. Quantitative RT–PCR confirmed a significant reduction in transcript abundance for MiPSK-θ and MiPSK-ω in dsRNA-treated juveniles relative to GFP-dsRNA controls (**Fig. 5A and 5B**). For MiPSK-α, transcript quantification in soaked pre-inoculation J2s was not informative, as both *MiPSK-α* and *MiPSK-δ* are expressed at very low basal levels in pre-parasitic J2s prior to host contact (**Fig. 4A, 4B**), rendering ΔΔCt-based quantification unfeasible at this stage. Both genes are strongly and specifically induced only upon host root attraction and early parasitism (**Fig. 4A, 4B**); thus, meaningful silencing is expected to occur after inoculation, once expression is triggered within the root. Tomato plants inoculated with *MiPSKs*-silenced juveniles developed significantly fewer galls than plants inoculated with control juveniles (**Fig. 5C and 5D**). Furthermore, the reproductive success of the nematodes was severely compromised, as evidenced by significant reduction in the number of egg masses per root system (**Fig. 5F**) and per gram of root tissue (**Fig. 5F**). These results demonstrate that reducing *MiPSKs* transcript abundance impairs feeding-site establishment and prevents development into reproductive females, consistent with a role for MiPSKs specifically in the post-invasion phases of parasitism.

**Figure 5.**
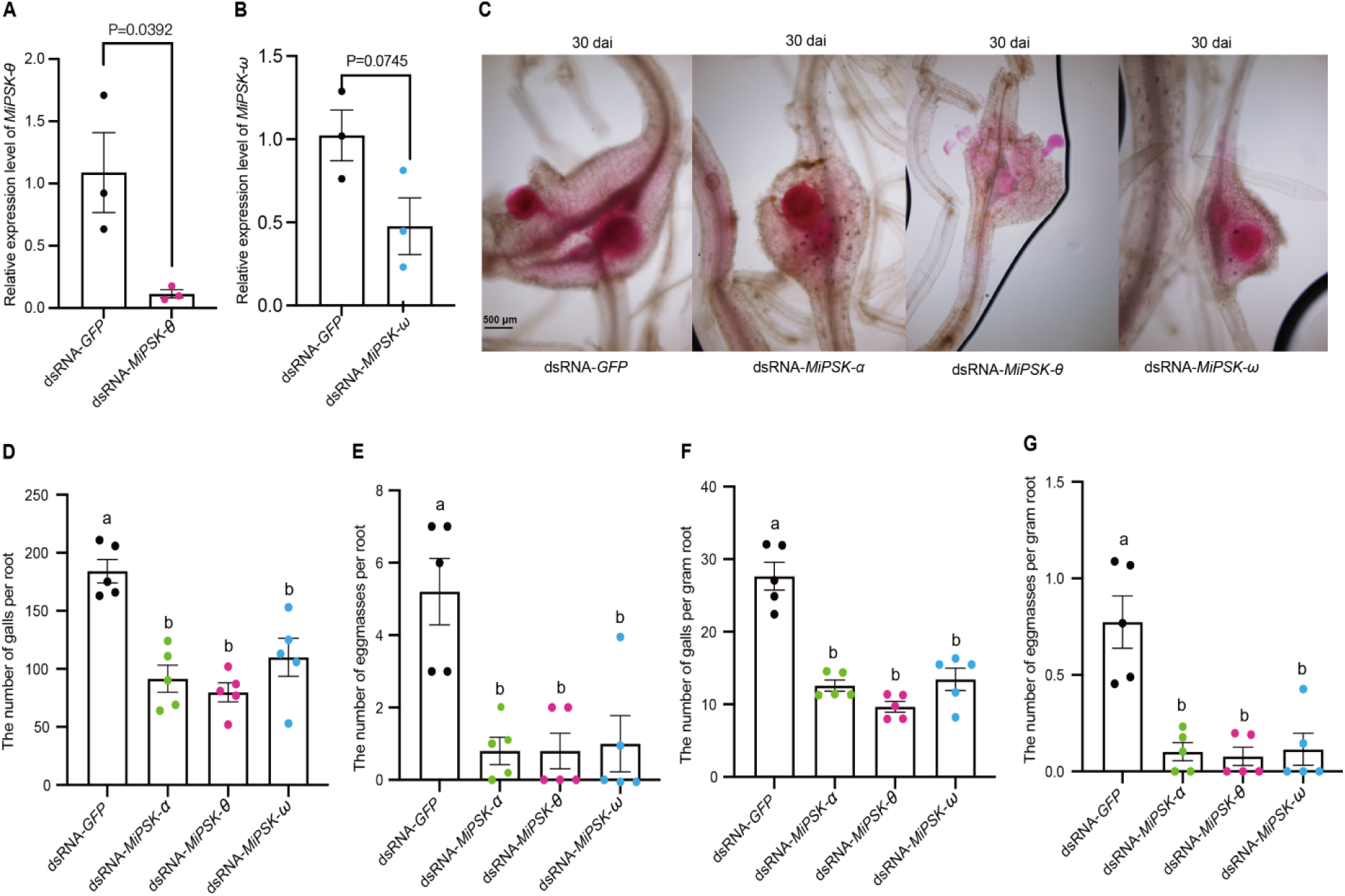
MiPSKs are required for successful parasitism of tomato by *M. incognita*. (**A, B**) Relative transcript abundance of MiPSK-θ (**A**) and MiPSK-ω (**B**) in pre-parasitic second-stage juveniles (J2s) following in vitro soaking in dsRNA targeting MiPSKs or eGFP (control). Transcript levels were normalized to 18S rRNA and quantified using the 2^−ΔΔCt^ method, with the *dsGFP*-treated group as the calibrator (set to 1). Bars represent the mean ± SEM of three technical replicates. Significant differences were determined using a two-sided Student’s t-test (P < 0.05). (**C**) Representative images of acid fuchsin-stained M. incognita infecting tomato roots following dsRNA-mediated gene silencing, at 30 days after inoculation (dai). (**D, E**) Numbers of galls (**D**) and egg masses (**E**) per root system at 30 dai. (**F, G**) Numbers of galls (**F**) and egg masses (**G**) per gram of root tissue at 30 dai. For panels (**D–G**), approximately 800 dsRNA-treated J2s were inoculated onto 14-day-old tomato seedlings, with six biological replicates per treatment. Statistical significance was assessed by one-way ANOVA followed by Tukey’s multiple range test. Different letters indicate statistically significant differences (P ≤ 0.05). Experiments were repeated two times independently with similar results; data from one representative experiment are shown. A third independent biological replicate is currently in progress and will be included in the final peer-reviewed manuscript. dsRNA, double-stranded RNA; J2, second-stage juvenile; eGFP, enhanced green fluorescent protein; dai, days after inoculation.

## Discussion

In this study, we report the identification and functional characterization of genes encoding phytosulfokine-like (PSK-like) peptides in plant-parasitic nematodes. To our knowledge, this represents the first report of a PSK mimic identified in any plant pathogen. Three converging lines of evidence establish these peptides as bona fide virulence effectors: First, *MiPSK* transcripts are highly induced during early infection and localize to the esophageal glands, the primary organelles for effector production and stylet-mediated delivery into host tissues. Second, the peptides are structurally competent for post-translational activation, encoding both the conserved DY sulfation site and, in the ω-class, a GGR amidation signal. Third, and most critically, RNAi-mediated knockdown of *MiPSKs* expression significantly impairs gall formation, reduces egg mass production, and prevents completion of the nematode life cycle, thus demonstrating that these peptides are functionally required for successful parasitism.

The phylogenetic distribution of nematode PSK-like genes offers some insight into the evolutionary origins of this mimicry strategy. Within the root-knot nematode lineage, PSK-like sequences are confined to Clade I *Meloidogyne* species, a pattern similar to PSY-like peptides (Yimer et al., 2023). However, PSK sequences were also identified in the cyst nematode *H. glycines* and the lesion nematode *P. vulnus*, pointing to a broader taxonomic distribution than was found for PSY mimicry. This suggests that PSK-based host manipulation may represent an ancient and phylogenetically widespread strategy among plant-parasitic nematodes. The apparent absence of both PSK- and PSY-like candidates across other PPN taxa should nonetheless be interpreted with caution, as fragmented genomic resources and the limited availability of high-quality long-read assemblies for non-model species may mask the presence of these short, repetitive, and structurally compact loci, which are particularly susceptible to assembly fragmentation and annotation gaps. Interestingly, while PSK loci were identified in the *P. vulnus* genome, they showed no detectable expression across different life stages examined (Dai et al., 2026b). These loci may therefore be induced only by specific host-derived cues or during transient infection windows not captured in available transcriptomes. Alternatively, they may represent evolutionary relics that have lost functional relevance in *P. vulnus*, which pursues a migratory rather than sedentary parasitic lifestyle and does not form the specialized feeding structures that PSK signaling is thought to promote. In contrast, the robust and stage-specific expression of *MiPSK* genes confirms that these peptides are actively deployed as pathogenicity factors at the onset of the host–parasite interaction in *M. incognita*.

To perform their functional role in plant developmental processes, PSKs require activation, as non-sulfated PSKs exhibit strongly reduced biological function (Kutschmar et al., 2009). Full PSK activity is dependent on tyrosine sulfation, which is catalyzed by tyrosyl protein sulfotransferases (TPSTs). Loss of TPST function in Arabidopsis results in dwarfism and root growth defects, underscoring the importance of PSK and related sulfated peptides in plant development (Komori et al., 2009). TPSTs are broadly conserved in eukaryotes, including nematodes (Igreja et al., 2022), suggesting that nematode PSKs may be sulfated prior to secretion. Such internal maturation would ensure that the peptides are in bioactive state, allowing for immediate interaction with the host receptors upon delivery via stylet. Additionally, PSK variants belonging to PSK-ω class terminate in conserved GGR motifs, which are canonical signals associated with propeptide processing and C-terminal amidation in nematodes (Li, 2018; Rosoff et al., 1993). This implies that these PSKs undergo sophisticated proteolytic cleavage and post-translational modifications to generate mature, bioactive ligands. Determining the precise *in vivo* biochemical status of these peptides, including their sulfation and amidation patterns, will be essential to resolve their functional diversity.

The presence of multiple PSKs arranged in clusters in root-knot nematode genomes suggests lineage-specific gene expansion, potentially allowing for functional specialization or different regulation during different infection phases. Given the broad host range of Clade I, which infects thousands of plant species, it is tempting to speculate that maintaining multiple PSK-like variants allows these nematodes to interact with divergent PSK receptor variants across host taxa, or to fine-tune signaling output at different infection stages. The tandem genomic arrangement of PSK-like loci may also facilitate dosage amplification, ensuring sufficient peptide production to saturate host PSK receptors during the narrow window of feeding site initiation.

In plants, PSK perception is mediated by leucine-rich repeat receptor-like kinases, primarily PSKR1 and PSKR2, which regulate growth, cell expansion, and stress responses (Matsubayashi et al., 2006; Stuhrwohldt et al., 2015; Amano et al., 2007; Zhang et al., 2018). Given that *pskr-1* mutants exhibit severe defects in feeding site development, it is plausible that nematode-derived PSKs engage these host receptors to activate developmental programs facilitating giant cell formation (Rodiuc et al., 2016). However, PSK signaling also attenuates pattern-triggered immunity, raising the alternative possibility that nematode-derived PSKs suppress early defense responses to facilitate root invasion (Hua et al., 2025). These two functions are not mutually exclusive and may operate at different time points during parasitism. Resolving this question will require receptor-binding assays with sulfated synthetic MiPSK peptides and genetic analysis in host lines with disrupted PSK receptor function.

Interestingly, nematodes PSK-like precursors differ in architecture from PSY-like mimics (Yimer et al., 2023). PSY-like mimics are highly compact, with the conserved PSY motif positioned immediately after the signal peptide, consistent with minimal processing beyond signal peptide cleavage and tyrosine sulfation. In contrast, PSK-like precursors contain extended and variable propeptide regions upstream of the conserved PSK motif, suggesting additional proteolytic modifications and potentially tighter regulatory control over peptide activation. These structural differences may also reflect distinct deployment strategies. The compact architecture of PSY-like precursors is consistent with secretion of near-mature peptides directly into the apoplast. By contrast, the extended propeptide regions of PSK-like precursors are more reminiscent of plant PSK precursors, which require subtilase-mediated cleavage for activation. This raises the intriguing possibility that nematode PSK-like peptides co-opt host subtilases for their own maturation — a strategy conceptually analogous to the exploitation of host SUMOylation and other post-translational machinery by nematode CLE mimics (Wang et al., 2021). Alternatively, propeptide processing may occur within nematode gland cells prior to secretion. Distinguishing these models will clarify whether the host cellular environment itself is required to generate bioactive MiPSK peptides.

In summary, this work establishes PSK peptide mimicry as a new layer in the molecular dialogue between plant-parasitic nematodes and their hosts. The identification of structurally and functionally distinct PSK-like classes in multiple nematode species suggests that exploitation of host PSK signaling is a conserved, and potentially adaptive, feature of sedentary endoparasitism. Future work should resolve the biochemical maturation of secreted MiPSK peptides, identify the host receptors and downstream signaling components they engage, and determine whether PSK mimicry operates primarily through growth promotion, immune suppression, or both. Beyond nematology, our findings raise the broader question of whether PSK mimicry is a virulence strategy shared with other plant-associated pathogens and symbionts that induce cell proliferation in host tissues.

## Material and Methods

### Identification and sequence analysis of nematode PSK genes

Nematode genes and transcripts analyzed in this study were identified using publicly available databases, including RKN genome resource (https://www6.inrae.fr/meloidogyne_incognita) and WormBase ParaSite (https://parasite.wormbase.org). Predicted translation products were analyzed for the presence of N-terminal signal peptides using SignalP-6.0 (https://services.healthtech.dtu.dk/service.php?SignalP). The presence of transmembrane domains was assessed using DeepTMHMM (https://doi.org/10.1101/2022.04.08.487609). Sequence logo analyses were performed using WebLogo (http://weblogo.berkeley.edu/).

### Phylogenetic analysis of PSK sequences

Phytosulfokine (PSK) peptide sequences from plants and nematodes were collected from publicly available genome and protein databases (**Supplementary Table 1**). Multiple sequence alignment was performed using MUSCLE (v3.8.1551) with default parameters. The resulting alignments were visually inspected and manually curated where necessary to minimize alignment errors, particularly in conserved regions. To eliminate poorly aligned and divergent regions, the alignment was further refined using trimAl (v1.4.rev15) with the automated1 mode, which selects the optimal trimming strategy based on the characteristics of the input alignment. Phylogenetic analysis was conducted using the maximum likelihood method implemented in IQ-TREE (v2.1.4-beta) with parameters “-m MFP -st AA -bb 1000 -alrt 1000 -nt AUTO -seed 2025”. The resulting phylogenetic trees were visualized and edited using iTOL.

### Plant growth condition and nematode infection assays

A single-egg mass population of *Meloidogyne incognita* was maintained on tomato plants (*Solanum lycopersicum* cv. Moneymaker) grown in sand under greenhouse conditions, as previously described (C. Wang et al., 2009). Eggs were collected from cultures maintained for approximately three months and subsequently incubated to obtain pre-parasitic second-stage juveniles (J2s) as described previously (Yimer et al., 2023). Tomato seeds (*Solanum lycopersicum* cv. Moneymaker) were germinated under controlled conditions for two weeks (26 °C; 16 h light/8 h dark). The resulting seedlings were transferred to individual pots, after which each plant was inoculated at the root zone with approximately 2,000 *M. incognita* J2s. The extent of nematode infection was evaluated at 30 days after inoculation (dai) as described previously (Yimer et al., 2023).

### RNA extraction, cDNA synthesis, and quantitative real-time PCR (qRT-PCR)

Nematode eggs, parasitic third- and fourth-stage juveniles (J3/J4), and adult females were collected from tomato roots infected for approximately three months. Eggs were incubated in a hatching chamber to obtain pre-parasitic second-stage juveniles (pre-J2s). To generate root-attracted J2s, tomato root tips were grown aseptically on Murashige and Skoog (MS) medium for two weeks and transferred to sterile plates. Pre-parasitic J2s were mixed with Pluronic F-127 gel (Sigma, USA) at a final concentration of 23% (w/v) and evenly spread over the plate surface to fully cover the root tips. After incubation at 28 °C in darkness for 6 h, nematodes accumulated around the root tips were collected and designated as root-attracted J2s (attracted-J2). Total RNA was isolated using the TransZol UP reagent (TransGen, China) following the manufacturer’s instructions. RNA concentration and purity were assessed using a NanoDrop OneC Microvolume UV–Vis spectrophotometer (Thermo Scientific, USA). First-strand cDNA synthesis was performed using the HiScript IV RT SuperMix for qPCR (+gDNA wiper) kit (Vazyme, China). Quantitative real-time PCR (qRT-PCR) was carried out using the iTaq Universal SYBR Green One-Step qRT-PCR Kit (Bio-Rad). Reactions were carried out in a total volume of 20 µL, consisting of 10 µL iTaq Universal SYBR Green reaction mix, 0.25 µL iScript reverse transcriptase, 1 µL of each primer (10 µM), 5.75 µL nuclease-free water, and 1.5 µL DNase-treated RNA (80 ng µL⁻¹). Amplification was performed on an Applied Biosystems QuantStudio 3 Real-Time PCR System (Life Technologies, USA) under the following cycling conditions: reverse transcription at 50 °C for 10 min, initial denaturation at 95 °C for 1 min, followed by 40 cycles of 95 °C for 15 s and 60 °C for 60 s. Melt curve analysis was performed by gradually increasing the temperature to 95 °C to confirm amplification specificity. Gene expression analyses were performed using two independent biological replicates, each analyzed with three technical replicates. Transcript levels of *MiPSKs* were normalized to the nematode reference gene *18S* rRNA and relative expression levels were calculated using the 2-ΔΔCT method (Livak and Schmittgen, 2001). The PCR primers are listed in **Table S1**.

### In situ hybridization

Total RNA was extracted from root-attracted J2s of *M. incognita* using the RNeasy Plant Mini Kit (Qiagen, USA), and first-strand cDNA was synthesized using the High-Capacity cDNA Reverse Transcription Kit (Applied Biosystems, USA). The full-length cDNA sequences of three *MiPSK* genes representing the α, θ, and ω classes were amplified by PCR and used as individual templates for probe synthesis. Primer sequences used for probe generation are listed in Table S1. Sense (negative control) and antisense single-stranded cDNA probes were synthesized independently for each gene by asymmetric PCR using a digoxigenin (DIG)-labeling mix (Roche, USA). The three antisense probes were subsequently pooled at equal concentrations prior to hybridization to enable simultaneous detection of transcripts across divergent *MiPSK* classes. Whole mount *in situ* hybridization was performed as described previously (Gao et al., 2003; Yimer et al., 2023). Nematodes were fixed in 4% paraformaldehyde in PBS buffer for 18 h at room temperature. The fixed nematodes were snap-frozen in liquid nitrogen and then crushed with a mortar and pestle to break the body wall, followed by permeabilization with proteinase K (0.5 mg mL⁻¹; Roche) at 37°C for 30 minutes. The nematodes were sequentially treated with methanol and acetone at −80 °C for 2 min each. After thorough washing, samples were pre-hybridized at 50 °C for 30 min, followed by overnight hybridization with DIG-labeled probes at 50 °C. Hybridized probes were detected using alkaline phosphatase–conjugated anti-DIG antibodies (Roche, USA). Images were captured using an Olympus IX71 microscope.

### Gene silencing

Double-stranded RNA (dsRNA) targeting the full-length coding sequence (CDS) of *MiPSKs* was synthesized as described previously (Dai et al., 2026b). Briefly, dsRNA was generated using the T7 High Yield RNA Transcription Kit following the manufacturer’s instructions (Vazyme, China). The T7 promoter sequence with protective nucleotides (5′-ggcTAATACGACTCACTATAGGG-3′) was appended to the 5′ ends of both forward and reverse primers (**Table S1**). dsRNA concentration and yield were quantified using a NanoDrop OneC Microvolume UV–Vis spectrophotometer (Thermo Scientific, USA).For RNA interference assays, approximately 10,000 freshly hatched pre-parasitic J2s were incubated in 200 µL soaking solution containing 1,000 ng µL⁻¹ dsRNA, 2 µL of 1% Triton X-100, and 2 µL of 1ng µL⁻¹ resorcinol. Nematodes were incubated for 50 h in darkness at room temperature with 15 rpm rotation, as described previously (Dai et al., 2026b).

Resorcinol was included to enhance dsRNA uptake. After incubation, J2s were washed five times with nuclease-free water and were divided into two aliquots, one aliquot was used to assess gene silencing efficiency by quantitative real-time PCR (qRT-PCR), while the other was used for plant infection assays. dsRNA targeting enhanced green fluorescent protein (*eGFP*) was used as a negative control. To evaluate silencing efficiency, total RNA was extracted from dsRNA-treated J2s, and *MiPSKs* transcript levels were quantified by one-step qRT-PCR as described above. For infection assays, tomato seedlings were germinated and grown in sand as described above. Two-week-old seedlings were inoculated with 800 dsRNA-treated J2s per plant. At 30 dai, the numbers of galls and egg masses per root system were recorded for six biological replicates per treatment, and fresh root weights were measured. Roots were subsequently stained with acid fuchsin to visualize nematodes, and the number of nematodes within root tissues was quantified under a light microscope. Statistical analyses and graphical visualization were performed using GraphPad Prism software version 9.5.1.

## Data and Materials Availability

All study data are included in the article and/or supporting information.

## Acknowledgments

This work was supported by the U.S. National Science Foundation under award IOS-2203286 and by the National Institutes of Health under grant 1R35GM158083-01, both awarded to Shahid Siddique.

## Supplementary Figures

**Figure S1.**
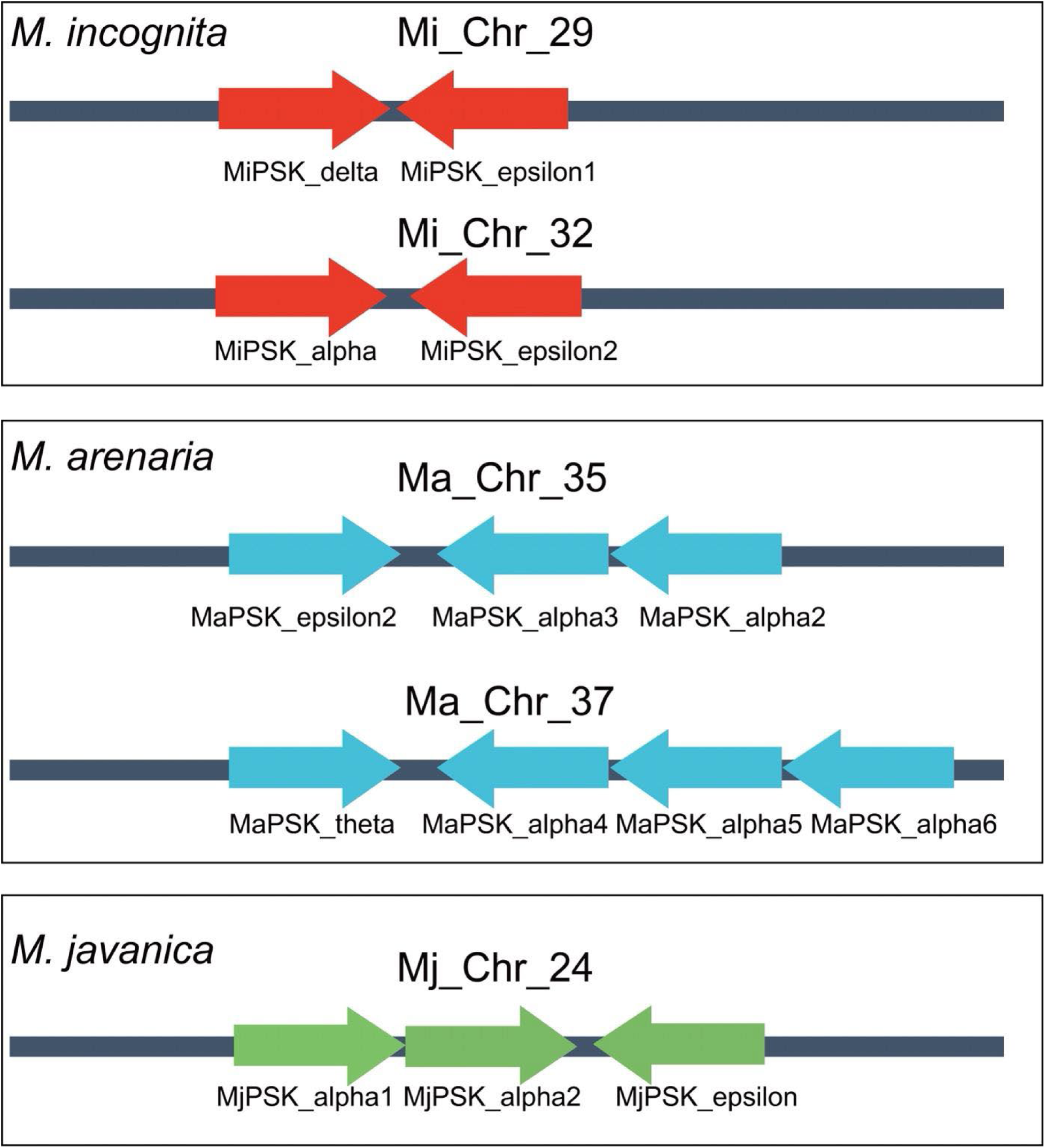
Genomic organization of PSK-like genes in *Meloidogyne incognita*, *Meloidogyne arenaria* and *Meloidogyne javanica*. PSK-like loci are clustered together in all three species. In *M. incognita*, four PSK-like genes are located on chromosome 29 and two on chromosome 32. In *M. arenaria*, three PSK-like genes (Ma_16833, Ma_16854, Ma_16855) are clustered on chromosome 35, and four genes (Ma_12898, Ma_12917, Ma_12918, Ma_12919) on chromosome 37. In *M. javanica*, three PSK-like genes (Mj_24829, Mj_24830, Mj_24854) are located on chromosome 24. Arrows indicate gene orientation; genes on the forward strand are shown pointing right and genes on the reverse strand pointing left.

